# Crosstalk Between Disrupted Blood-Brain Barrier, Neuroinflammation, and Coagulopathy in AC70 hACE2 Tg Mice with Prolonged Neurological Manifestations Induced by the Omicron BA.1 Variant of SARS-CoV-2

**DOI:** 10.1101/2025.04.03.646981

**Authors:** Aleksandra K. Drelich, Panatda Saenkham-Huntsinger, Yu-Hsiu Wang, Xuping Xie, Barbara M. Judy, Thomas G. Ksiazek, Bi-Hung Peng, Chien-Te K. Tseng

## Abstract

Long COVID, or post-acute sequela of COVID (PASC), has emerged as a major post-pandemic challenge, with lasting neurological effects in a substantial number of patients. The underlying pathophysiological mechanisms of PASC remains elusive, largely due to the lack of suitable animal models that replicate key clinical and pathological features, especially in survivors of acute infection. Here, we employed the Omicron BA.1-infected AC70 hACE2 Tg mice to explore the pathogenesis of COVID-19. Our findings in the AC70/BA.1 model demonstrated that surviving mice developed lasting neurological sequelae of COVID-19, making it a promising platform for studying the pathogenesis of neuro- or long-COVID. Histopathological examinations revealed a transient pulmonary inflammatory response that correlated with the levels of live virus, whereas neuroinflammation within the brains persisted and progressed beyond the acute phase, without a clear linkage to direct virus effects. We showed that the severity of neurological sequelae directly correlated with the degree of blood-brain-barrier (BBB) disruption, alterations in tight junctions (TJs) and fibrinogen extravasation into the brain parenchyma. Additionally, BA.1-infected mice elicited a sustained systemic prothrombotic state, with dysregulated coagulation and fibrinolytic pathways in an organ-specific manner, as evidenced by distinct pattern of intracellular fibrin(ogen) depositions, elevated d-dimers, and tissue-plasminogen activator (t-PA) expression in the lungs and brains. We also found a positive correlation between progressive neuroinflammation and t-PA expression, which was closely co-localized with Iba-1-positive cells, indicating that activated microglia may serve as an additional source of t-PA in the CNS. Lastly, we showed that BA.1 variant triggered prolonged anti-Annexin A2 (ANXA2) autoantibody production. ANXA2 is essential for the neurovascular system, coordinating multiply dynamic processes within the endothelium, including cell surface fibrinolysis and TJ assembly. Our analysis revealed that impaired TJ structures coexisted with diminished ANXA2 levels around the brain vessels, suggesting its involvement in BBB dysfunction and neuroinflammation.

**Author Summary:** Long COVID symptoms, especially neurological complications, remains a key public health concern worldwide. Alterations in the cerebrovascular system, along with ongoing coagulation disorders, may play a crucial role in the pathogenesis of this condition. Here, we established murine model for study post-acute sequela of COVID (PASC). We demonstrated that AC70 Tg mice infected with a high dose of BA.1 Omicron variant developed prolonged neurological symptoms, accompanied by progressive neuroinflammation in survivors. Our findings clearly showed the involvement of blood-brain-barrier (BBB) disfunction in the severity of neuropathology induced by SARS-CoV-2 infection. We also showed that the bidirectional interaction between dysregulated coagulation/ fibrinolysis and inflammatory systems may exhibit a distinct pattern in the CNC, significantly contributing to persistent neurological symptoms. Finally, we highlighted a potential role of malfunctioning Annexin A2 (ANXA2) in this phenomenon that may serve as a promising drug target for neuro-COVID interventions.

## Introduction

Over four years after the onset of the COVID-19 pandemic, millions of people continue to suffer from the consequences of the infection with new variants of concern (VOCs), e.g. Omicron and its sub-lineages, that escape from natural and vaccine-induced immunity, as well as long-term sequela after recovery from the initial infection, referred to as long-COVID syndrome or post-acute sequelae of COVID (PASC). COVID-19 was originally described as severe respiratory syndrome limited to the respiratory tract. However, multiple retrospective cohort studies of COVID-19 survivors indicate that the virus may impact multiple organs, with a significant effect on the Central Nervous System (CNS). Although the prevalence may vary, studies show that up to 80% of hospitalized cases may experience neurological signs during the acute phase of disease [**1,2**], with some persisting for over six months [**3,4**]. Frequently reported manifestations include headache, anosmia, cognitive dysfunction, cerebrovascular events, encephalopathy, neuropathies, strokes, seizures, and myopathy [**5–10**]. Some reports described abnormal movements, syncope, psychomotor agitation, or autonomic dysfunction [**11, 12**].

While the exact mechanisms behind SARS-CoV-2 neurovirulence remain unclear, postmortem studies of patients who succumbed to COVID-19 have revealed widespread inflammatory responses in the CNS, supporting the role of neuroinflammation in neuro-COVID. Multifocal glial cell activation, particularly microglia, in various brain regions [**13–16**] are a common finding in COVID patients. Studies in a mouse model of mild COVID-19 revealed that activated microglia persisted for up to seven weeks after infection with prototype isolate of WA-1, suggesting that microglia could be partly responsible for neurological PASC [**17**]. However, the exact mechanism of neuroinflammation and the role of activated microglia and other CNS resident cells in host responses during COVID has been poorly defined.

Possible mechanisms contributing to SARS-CoV-2-related neurological damage and CNS inflammation include direct viral invasion of the CNS, as well as indirect effects driven by systemic inflammatory dysregulation, autoimmunity, and/ or coagulation dysfunction, potentially leading to increased vascular permeability or the blood-brain-barrier (BBB) disruption.

The ability of SARS-CoV-2 to directly access the CNS, either through peripheral nerves and/or via the hematogenous route, along with its potential to establish a productive infection in brain, has been thoroughly investigated. Inconsistent findings on the direct infection of endothelial cells (ECs), and lack of clear evidence for productive infection of the brain parenchyma *in vivo* [**18–21**], suggest that alternative mechanism(s) are likely involved.

COVID-associated coagulopathy (CAC) is complex syndrome characterized by imbalanced coagulation-fibrinolysis pathways that complicates acute phase of SARS-CoV-2 infection, most frequently manifested by pulmonary embolism at the early stage of infection. The persistence of systemic coagulopathy may lead to cerebrovascular alterations and extracellular and intracellular fibrin/ fibrinogen deposits in brain of COVID patients, suggesting the involvement of CAC in the development of neurological complications. Cerebral homeostasis and proper neuronal function are regulated by the BBB, a specialized endothelial structure within cerebral blood vessels that selectively restricts the paracellular diffusion of compounds from the bloodstream into the brain. The barrier properties of these ECs rely heavily on tight junctions (TJs), which form connections between neighboring cells. Sustained systemic coagulopathy may impair BBB function, leading to subsequent extravasation of immune cells and serum components, such as IgG, fibrinogen and other blood-borne thrombo-immunomodulators, into the brain parenchyma, thereby promoting neuroinflammation and coagulation cascades. A notable finding in COVID-19 patients experiencing neurological disorders is the detection of cerebral vascular dysfunction, where hemorrhages and ischemia are common features of CNS pathology [**22–24**]. Other studies on deceased patients have clearly demonstrated cerebrovascular injury, characterized by the extravasation of hemostatic components, platelet aggregation, and abnormal coagulation [**13**], suggesting BBB disruption or leakage. A recent report highlights a potential correlation between BBB impairment and cognitive dysfunction, observed during acute infection and post-acute phase of disease [**25**]. Collectively, an expanding body of evidence underscores vascular impairment and BBB disruption as a possible mechanism underlying the connection between systemic coagulopathy and neuroinflammation.

Understanding the pathophysiological basis of this mechanism is essential for gaining deeper insights into the neurological effects of COVID-19. We have recently reported that AC70 hACE2 Tg mouse model infected with USA-WA-1 isolate recapitulates several unique features of CAC. Notably, we demonstrated that SARS-CoV-2 affects the Annexin A2 (ANXA2) system in our mice and proposed that virus-induced anti-ANXA2 autoantibodies may contribute to fibrinolysis resistance [**26**], a phenomenon commonly seen in CAC [**27**]. ANXA2 is a multi-compartmental protein that exists either as a monomer or forms a heterotetrameric complexes (AIIt) with the associated small S100A10 (P11) proteins on the EC surface. Extracellular ANXA2, especially in AIIt complexes, plays a crucial role in the neurovascular system by orchestrating multiply dynamic processes within the endothelium. By serving as a co-receptor for plasminogen and tissue plasminogen activator (t-PA), key factors in plasmin generation, the AIIt complexes promotes cell surface fibrinolysis, thereby regulating fibrin balance, an essential process for maintaining vascular homeostasis in the brain. Furthermore, ANXA2, within the AIIt complex, has been identified as an endothelial F-actin-binding protein involved in junction assembly [**28**], playing a pivotal role in TJ integrity, vascular permeability control, and inflammatory modulation [**29, 30**]. Consequently, dysregulation of ANXA2 system may be an important mechanism behind COVID-related cerebrovascular disorders.

Yet, studies relying on human samples face limitations, including inconsistent collection timing that hinder early infection analysis before symptom onset, the heterogeneity of patient cohorts, and sample sources primarily obtained from outside the CNS or cerebrospinal fluid (CSF). Thus, establishing a well-representative animal model that captures multiple facets of neuro-COVID, particularly in its post-acute stage, remains a pressing need. In this study, we leveraged the more recent Omicron variant of SARS-CoV-2 to evaluate its pathogenesis in AC70 hACE2 Tg mice. We demonstrated that AC70/BA.1 model exhibited prolonged neurological sequelae beyond the acute infection phase, enhancing its suitability for PASC studies. Importantly, our results convincingly indicate a positive correlation between the severity of neurological sequelae, neuroinflammation and the extent of BBB disruption, occurring in conjunction with tissue/ organ-specific coagulation dysfunctions. Finally, we highlighted the potential role of ANXA2 system in this phenomenon.

## Results

### Physical and neurological impacts of SARS-CoV-2 Omicron BA.1 variant infection in AC70 hACE2 Tg mice

We have previously shown that AC70 hACE2 Tg mice infected with high dose (≥10^3^ TCID_50_) of original US-WA1/2020 isolate of SARS-CoV-2 resulted in 100% mortality within 5 days post-infection (dpi) [**26, 31**]. Here we demonstrate that high dose of Omicron variant causes a milder infection and disease, with delayed clinical symptoms compared to the original strain. Briefly, a cohort of 20 mice was challenged with ∼2.3×10^4^ TCID_50_, of the BA.1 variant (strain: EHC_C19_2811c) of SARS-CoV-2 and monitored for up to 21 dpi. We noticed that infected mice developed attenuated clinical disease and displayed a pattern of biphasic course. During the early phase, between 4-5 dpi, mice showed mild symptoms with slight weight loss. In the later phase, between 7-9 dpi, ∼64.3 % infected mice exhibited moderate to severe illness characterized by an average of ∼ 14 ± 10 % weight loss, which typically peaked at 9 dpi. Mortality reached up to ∼20% between 8–9 dpi in repeated studies. The surviving infected mice displayed clinical signs, including ruffled fur, lethargy, hunched posture, and orbital tightening but gradually recovered by 11 dpi and remained alive through 21 dpi, when the studies were terminated [**Figure 1A-C**]. Interestingly, we observed for the first time the onset of severe neurological sequelae in ∼27% of BA.1-infected survivors, manifested by gait disorder, ataxia, as well as varying degrees of brachial or crural monoplegia and fore- or hind-leg paralysis, that typically occurred between 9 to 13 dpi and persisted through 21 dpi after the initial illness had fully resolved. [**Figure 1D**]. In this manuscript, the recovery phase, marked by the regaining of lost body weight in mice, is considered as the post-acute stage of the disease [**Figure 1A&B**].

**Fig 1.**
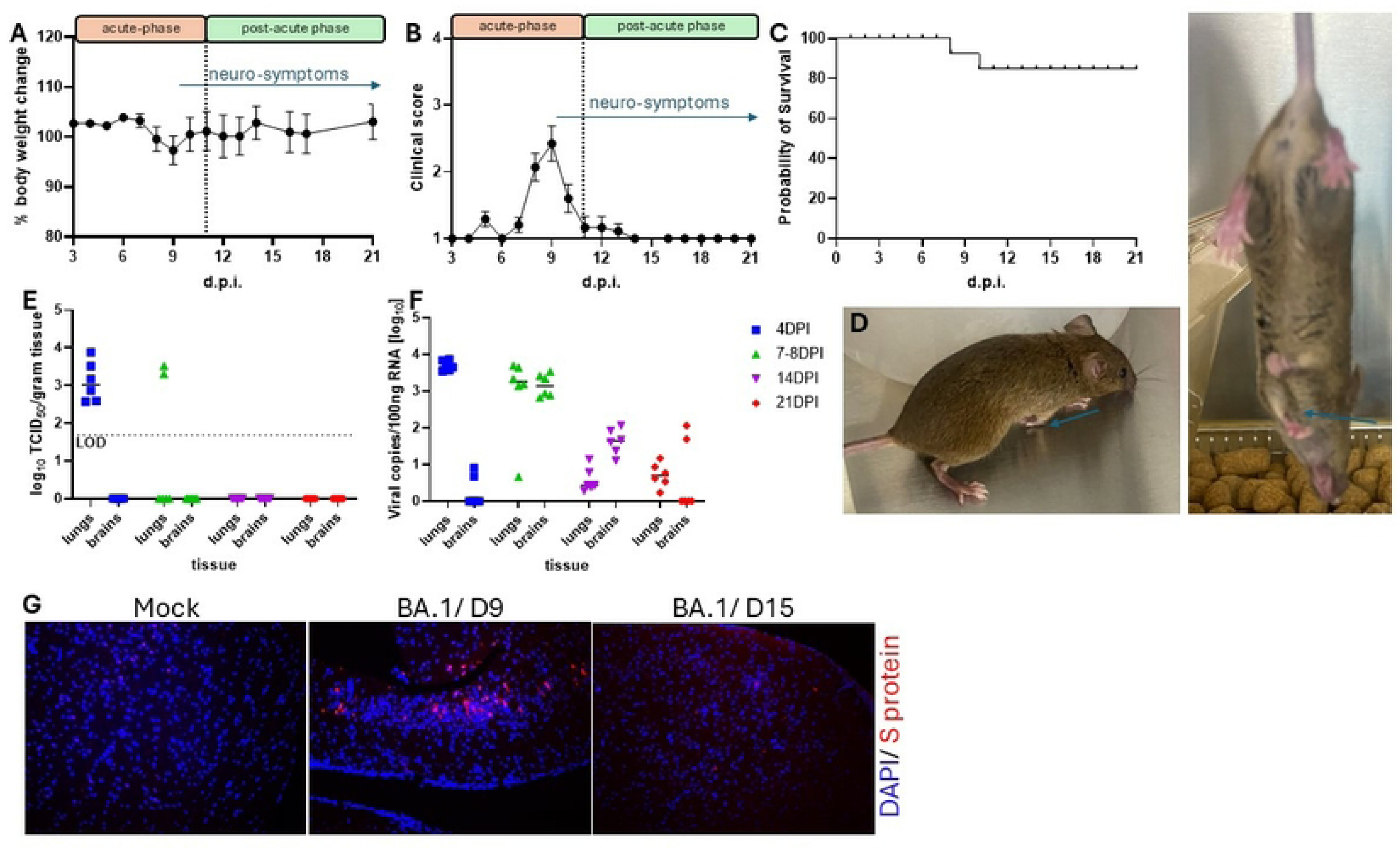
AC70 hACE2 Tg mice challenged with a high dose of Omicron BA.1 variant of SARS-CoV-2 resulted in attenuated infection and prolonged neurological manifestations. AC70 hACE2 Tg (N=20) were intranasally (i.n.) challenged with ∼2.3×10^4^ TCID_50_ of BA.1 (strain: EHC_C19_2811c) of SARS-CoV-2. Challenged mice were monitored daily for morbidity (weight loss and other clinical signs) and mortality. (A) Average percentage (%) of daily body weight changes after viral infection, expressed as Mean ± SEM (standard error mean) were plotted. (B) Clinical signs rated on a 4-point (0-4) scale. (C) Accumulative mortality rates. (D) Representative photographs depicting severe neurological sequelae in surviving mice, manifested by right upper limb paralysis. Tissues/organs (N=6 per time point, t.p.) were harvested from infected mice at 4-, 7-, 14- and 21- dpi. (E) Yields of infectious progeny virus in lungs and brains were determined using a standard Vero E6 cell-based infectivity assay and expressed as log_10_ TCID_50_ virus per gram of tissue. The dotted horizontal line indicates the limit of detection (LOD, ∼1.69 log_10_ TCID_50_/gm). (F) Total RNA extracted from tissue samples was analyzed using qRT-PCR assays targeting the SARS-CoV-2 E gene to determine viral RNA levels. (G) Representative immunofluorescence (IF) staining of brain sections at 9 and 15 dpi. Mock slides were included as controls. The images depict staining for nuclei (DAPI, blue) and SARS-CoV-2 Spike (S) antigen (red). Scale bar: 100 µm. Two independent studies were performed.

### Kinetics of Omicron BA.1 replication within the lungs and brains of AC70 hACE2 Tg mice

Lung and brain are two prime targets of zoonotic human respiratory SARS-CoV, SARS-CoV-2 and MERS-CoV infection in established hACE2- and hDPP4-Tg mice, respectively [**26, 32, 33**]. We explored the kinetics and magnitude of the less pathogenic Omicron BA. 1 variant infection within these two organs. Briefly, we intranasally challenged a group of 36 AC70 hACE2 Tg mice and euthanized six mice at each of the following time points 4, 7, 14, and 21 dpi. Tissue specimens were harvested and processed to determine live virus yields and viral RNA using a standard Vero E6-based infectivity and qRT-PCR assays, respectively. We showed that an average of ∼ 3.1 ± 0.5 and 3.4 ± 1.8 log_10_ TCID_50_/per gram of infectious virus could be recovered from the lung homogenates of all six and two out of six mice at 4 and 7 dpi, respectively. No detectable live virus was present in the lungs at 14 and 21 dpi. We were unable to recover any infectious virus from the brains throughout all the time points tested [**Figure 1E**]. Next, we performed semi-quantitative RT-PCR analysis to further characterize the viral RNA kinetics within these two main targets of SARS-CoV-2 infection. As shown in **Figure 1F**, we observed a gradual decline in viral copies in lungs across each group of six mice, with average of ∼ 3.7, ∼ 3.2, to < 1.0 log_10_/per 100 ng total RNA at 4-, 7-, 14-, and 21-dpi, respectively. While no live virus was detected in the brains over time, viral RNA was readily detected in some mice tested. Specifically, we noted that 2 brains, out of 6, harvested at 4- and 21-dpi, respectively, possessed lower but detectable levels of viral genome, whereas all 6 brains assessed at 7 and 14 dpi had higher levels of viral genome, especially at 7 dpi.

Given the noticeable presence of viral RNA in the brains between 7 and 14 dpi, we applied standard indirect immunofluorescent (IIF) staining to determine whether the viral Spike (S) protein was localized within the brains of mice displaying neurological symptoms. We were able to detect patches of S-positive (S^+^) neuronal cells in the cortex and, to a lesser extent, the thalamus of all three mice tested at 9 dpi. The S^+^ neuronal cells remained sporadically detectable within the cortex at 15 dpi, but not at 21 dpi [**Figure 1G**].

### Histopathological changes in lungs and brains of Omicron BA.1-challenged AC70 hACE2 Tg mice

To investigate the impact of Omicron BA.1 infection on the host response, we evaluated the histopathology of lungs and brains of infected mice at 7, 9, 14 and 21 dpi. In lungs, we found that foci of inflammatory mononuclear cells could be unambiguously observed at 7 dpi, mainly within perivascular/peribronchiolar regions and sporadically around septal capillaries [**Figure 2A**]. We also noted a prominent blood clot formed within a pulmonary arteriole [**Figure 2B**], suggesting an onset of coagulation disorders. The extent of cellular infiltration within the lungs gradually subsided thereafter. Specifically, pulmonary inflammatory responses, as evidenced by the formation of perivascular cuffing, could only be sporadically detected at 9 dpi [**Figure C&D**], but not 14 and 21 dpi [Data not shown] in mice with or without suffering from severe neurological disorders. These results suggest to us that Omicron BA.1 infection induced transient inflammatory responses within the lungs of AC70 hACE-2 Tg mice.

**Fig 2.**
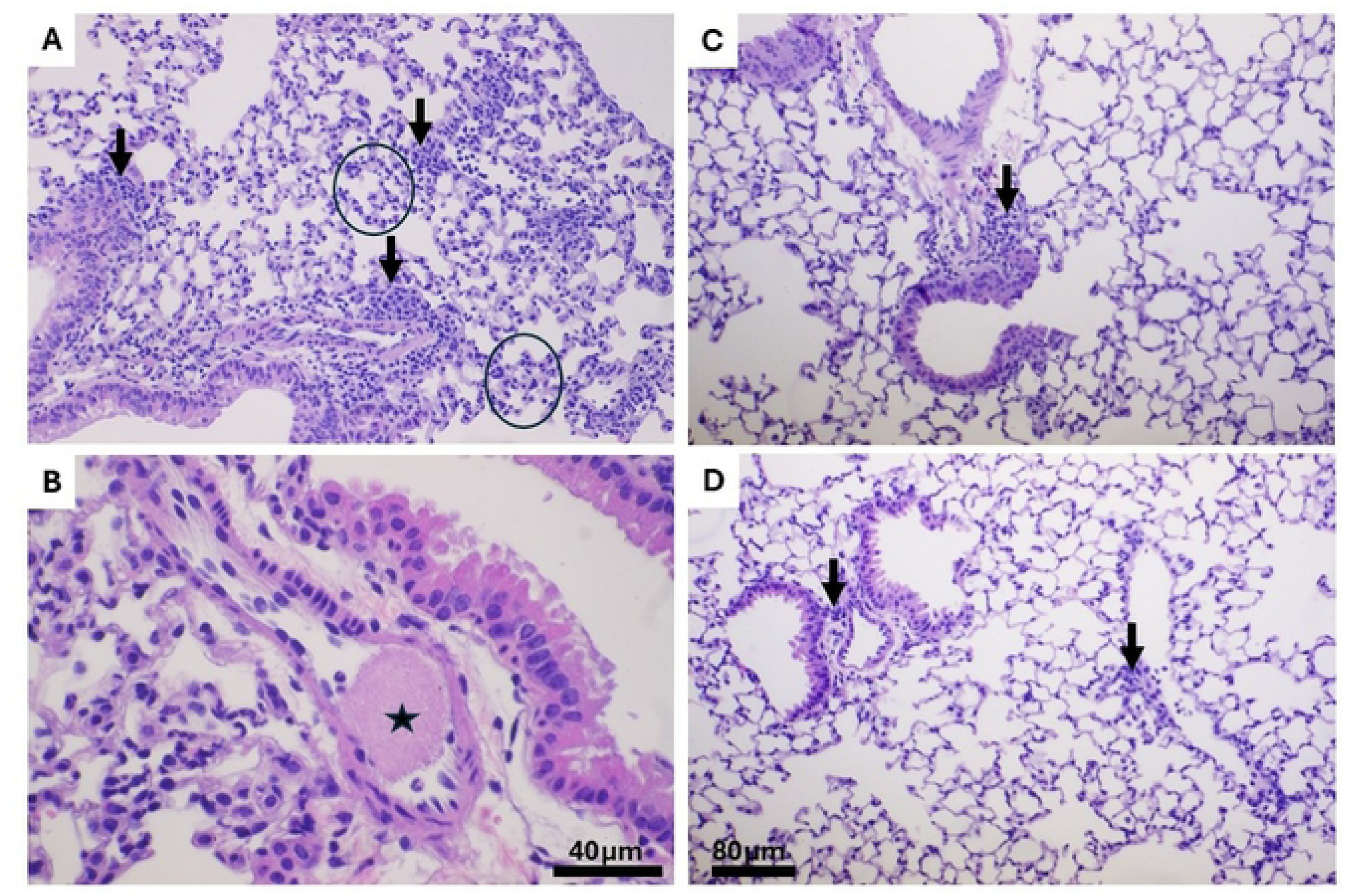
Histopathological examination of the lungs from BA.1-infected AC70 hACE2 Tg mice revealed transient inflammation and pulmonary embolic events. Lung tissue specimens (N=4 per group) harvested at 7-, the onset of clinical signs, and 9-dpi., the onset of neurological sequelae, from AC70 hACE2 Tg mice challenged with ∼2.3×10^4^ TCID_50_ of BA.1 of SARS-CoV-2 were subjected to the histopathology examination. Briefly, paraffin-embedded, and formalin-fixed tissue sections were stained with hematoxylin and eosin (H&E), as described in Materials and Methods. At 7 dpi, mononuclear cell infiltrations had extended from perivascular/peribronchiolar regions (arrows in **A**) to septal capillaries (**circles**). By 9 dpi, only few foci of perivascular cuffing left in mice that developed (arrow, **D**), and without (arrow, **C**) severe neurological symptoms. Blood clot (star in **B**) observed in a pulmonary arteriole. Scale bar = 80µm for A, C & D; 40µm for B. Representative images from two independent experiments.

Histopathological evaluation of the brains of Omicron BA.1-infected mice over time revealed noticeable mononuclear cell infiltrates in the cortex, meninges, and brainstem at 7 dpi [**Figure 3A**]. Neuroinflammation, accompanied by microglial activation, exacerbated at 9 dpi, when mice started developing neurological symptoms [**Figure 3B**, **3E&F**]. Interestingly, disrupted cerebellar white matter tract [**Figure 3D**] and deaths of neuronal cells that were identified as “red” neurons or apoptotic bodies [**Figure 3E&F**] could also be detected in those mice suffered from severe neurological symptoms, but not those with mild sequelae [**Figure 3C**]. Neuroinflammatory responses, mainly characterized by the presence of microglial activation and perivascular cuffing within deep cerebellar nuclei and brainstem, persisted at 14 dpi in mice suffered from either mild or severe neurological disorders [**Supplemented Figure 1A&B**]. However, such characteristic neuroinflammatory responses could be further extended to other brain regions, including meninges and neocortex at 21 dpi only in mice suffered from severe, but not mild, neurological disorders **[Supplemented Figure 1C&D]**.

**Fig 3.**
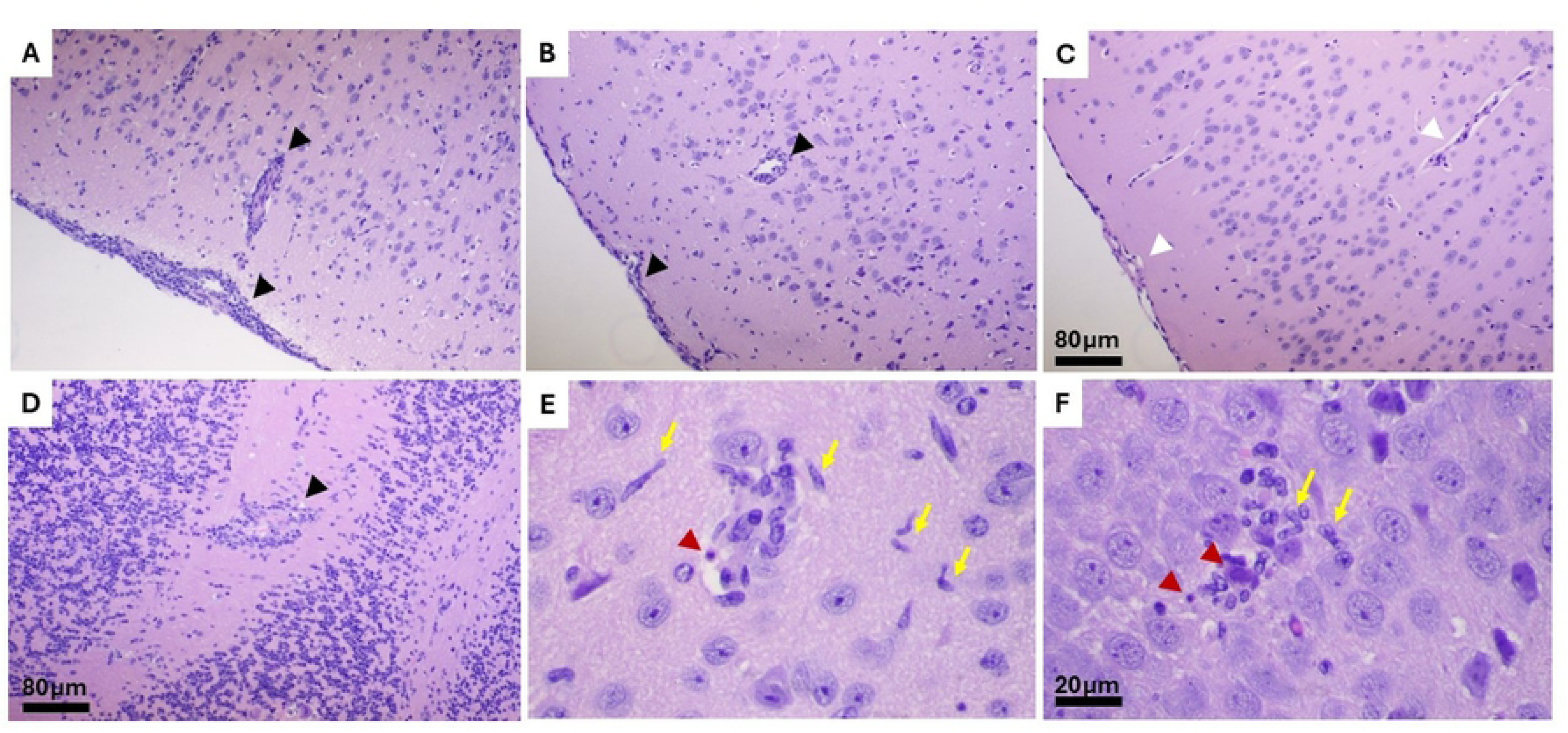
Histopathological examination of the brains from BA.1-infected AC70 hACE2 Tg mice revealed progressive neuroinflammation proportional to the severity of neurological manifestations. Brain tissue specimens (N=4 per group) harvested at 7-, the onset of clinical signs, and 9-dpi., the onset of neurological sequelae, from AC70 hACE2 Tg mice challenged with ∼2.3×10^4^ TCID_50_ of BA.1 of SARS-CoV-2 were subjected to the histopathology study. **M**ononuclear infiltrations were seen in the cortex, meninges (arrowheads in **A**), and brainstem in 3 of 4 mice in 7 dpi. By 9 dpi (**B**), more areas were involved, and the infiltrated cells migrated to surrounding where activated microglia were also present (yellow arrows in **E** and **F**). Interruption of white matter tract (cerebellar white matter in **D**), or neural cell death were seen as red neurons and apoptotic bodies (red arrowheads in **E** and **F**). No detectable pathology was seen in the brain (**C**) of mouse without severe neurologic symptom at 9 dpi. Scale bar = 80µm for A to D; Bar = 20µm for E and F. Representative images from two independent experiments.

To further validate glial cell activation, we performed the standard IHC and immunofluorescent staining assays by using specific antibodies against Ionized calcium-binding adapter molecule-1 (Iba-1) and glial fibrillary acidic protein (GFAP) antibodies, specific markers for microglial cells and astrocytes, respectively, for those mice suffered from severe neurological disorders. As shown in **Figure 4**, activated microglia (Iba-1^+^) were readily detectable at both 14 and 21 dpi, with more prominent presence at 21 dpi, along with distinct activation of astrocytes (GFAP+). Taken together, these results suggest that neuroinflammatory response triggered by Omicron BA.1 infection is likely an ongoing pathological process, starting from 7 dpi through at least 21 dpi.

**Fig 4.**
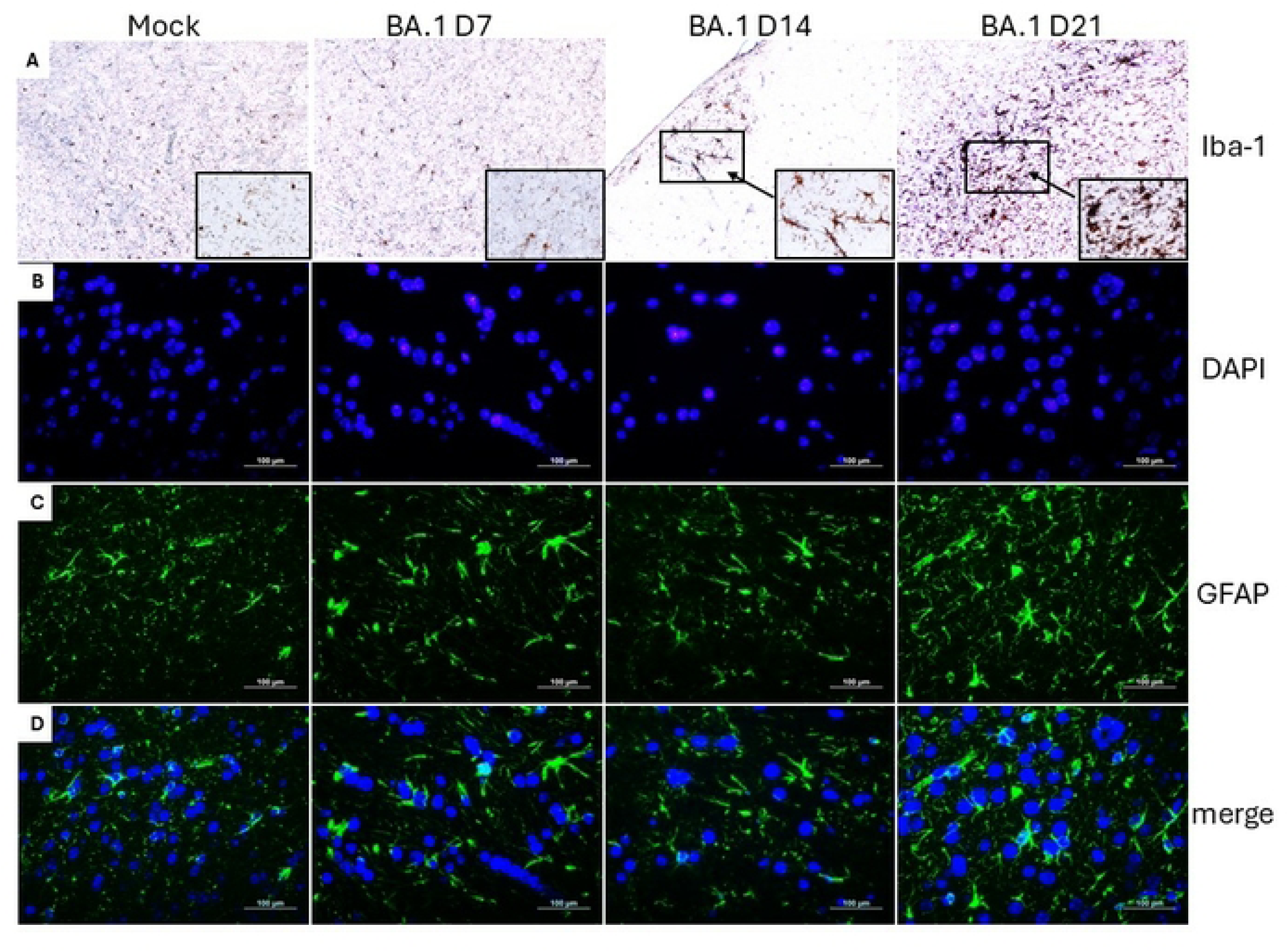
BA.1 variant of SARS-CoV-2 induced glial cell activation in AC70 hACE2 Tg mice. Brain tissue specimens harvested at 7-, 14- and 21- dpi. from AC70 hACE2 Tg^+^ mice challenged with ∼2.3×10^4^ TCID_50_ of BA.1 of SARS-CoV-2 were subjected to the immunostaining analysis. Mock controls were included. (**A**) Immunohistochemistry (IHC) indicating microglia cell activation. Microglia cells (brown) were stained with anti-Iba-1 antibody. Higher magnification views of the boxed sections are displayed for each time point. Scale bar= 100 µm. (**B-D**) Immunofluorescence (IF) staining of astrocyte activation. Astrocyte (green) were stained with anti-GFAP antibody. DAPI (blue) was used to show nuclei. The bottom panels display the merged images. Scale bar= 100 µm. Representative images from two independent experiments.

### Omicron BA.1 variant infection led to organ-specific coagulation disorders in AC70 hACE2 Tg mice

We have previously shown that intranasal infection with a high dose (> 10^4^ LD_50_) of SARS-CoV-2/WA1 isolate of AC70 hACE2 Tg mice resulted in the onset of key characteristics of COVID-associated coagulopathy (CAC) [**26**]. While coagulopathy is a hallmark of acute SARS-CoV-2 infection, multiple studies have demonstrated that coagulation abnormalities can persist well beyond the resolution of the acute phase, thereby suggesting a potential linkage between imbalanced hemostasis and the development of neuro- or long-COVID. Utilizing the Omicron BA.1/AC70 hACE2-Tg mouse model, we monitored the temporal dynamics of circulating d-dimer levels, a degradation product of cross-linked fibrin clot, in mice that survived SARS-CoV-2 infection. We found that the average of d-dimer levels in the circulation of 5 mice measured at each of indicated time points were significantly increased from the base value of 2402 ± 262 ng/mL measured before viral infection to 4217 *±* 174 (p<0.0001), 3763 *±* 214 (p=0.0009), 4063 *±* 208 (p<0.0001) and 4063 ± 208 ng/mL (p<0.0001) assessed at 4, 7, 14 and 21 dpi, respectively [**Figure 5A**] indicating a systemic prothrombotic state has been induced. As lung and brain are two prime pathological targets of SARS-CoV-2 infection, we investigated the contents of d-dimer within these two tissues/organs harvested between 9-10 dpi and at 21 dpi from mice suffered from neurological disorders. Similar to the rise observed in circulation, d-dimer content was also elevated in both organs, although the expression kinetics varied. Specifically, levels of d-dimer in the lungs were transiently increased from 1441 ± 103 ng/mL prior to infection to 5741 ± 1797 ng/mL at 9-10 dpi (p=0.0173) and subsequently returned to the level of ∼ 1789 ± 298 ng/mL at 21 dpi, a value not significantly different from the background [**Figure 5B**]. In contrast, we found that the levels of d-dimer within the brains increased from the basal value of 178.6 ± 36 ng/mL to 520 ± 45 ng/mL, albeit statistically insignificance, at 9-10 dpi, and continuously increased to 605.7 ± 114 ng/mL (p=0.02) at 21 dpi [**Figure 5C**]. As the contents of d-dimers often reflect the state of severity of thrombosis and/ or tissue damage, it is conceivable that the induction, intensity, and kinetics of coagulopathy triggered by SARS-CoV-2 infection might be tissue- or organ-specific.

**Fig 5.**
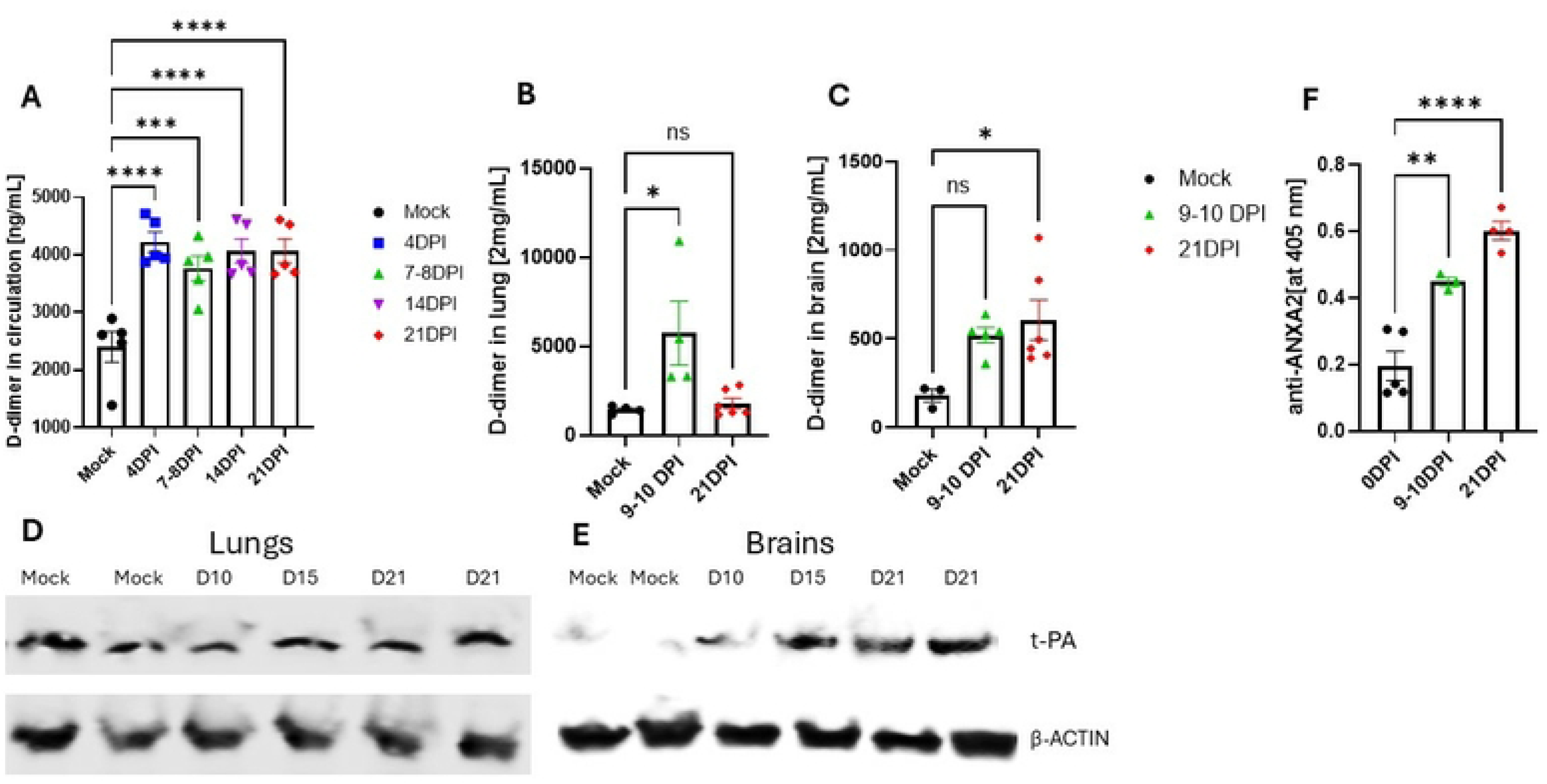
BA.1 variant of SARS-CoV-2 triggered coagulation disorders in an organ-specific manner in AC70 hACE2 Tg mice. (**A**) The concentration [ng/mL] of circulating D-dimers detected by ELISA-based analysis in gamma irradiated EDTA-plasma specimens derived from uninfected control and virally challenged mice at 4-, 7/8-, 14- and 21- dpi. with ∼2.3×10^4^ TCID_50_ of BA.1 of SARS-CoV-2 (n=5 per group). (**B-C**) The concentration of tissue D-dimers at 9/10- and 21 dpi. in (**B**) lungs and (**C**) brains. (**D-E**) The expression of tissue-plasminogen activator (t-PA) in (**D**) lungs and (**E**) brains measured by Western Blot (WB). β-actin was used as loading controls. Representative images from two independent experiments. (**F**) The levels of anti-Annexin A2 (ANXA2) in gamma irradiated EDTA-plasma samples analyzed by ELISA assay. The results were expressed as absorbance values at 405 nm.

Tissue plasminogen activator (t-PA) is a major player of the fibrinolytic process that breaks down blood clots to maintain a balance between blood clotting formation and dissolution. Therefore, we also investigated the expression of t-PA within the lungs and brains by using Western blotting-based analysis to examine whether the fibrinolytic pathways could be affected upon BA.1 infection. As shown in **Figure 5D&E**, we were only able to identify an increased trend of t-PA expression within the brains, but not in the lungs. Collectively, these findings indicate that fibrinolysis may be regulated in distinct ways within these two organs. Endothelial cells lining the blood vessels have been recognized as the prime source of t-PA [**34, 35**]. However, glial cells and other brain cellular components have been reported as potential sources of t-PA in the CNS [**36, 37**]. As profound microglial activation has been detected in Omicron BA.1-infected mice [**Figure 4]**, we investigated whether they could contribute to elevated t-PA expression in the brain by using double immunostaining against t-PA and Iba-1, respectively. As shown in **Figure 6**, consistent with the data of Western blotting analysis (**Figure 5D-E**), we also found an increased t-PA expression within the brain at 15 dpi, compared to the mock-infected control. Most importantly, the elevated expression of t-PA was closely associated or co-localized with activated microglial cells (Iba-1^+^), implying that these cells could serve as an additional source of t-PA in infected brains.

**Fig 6.**
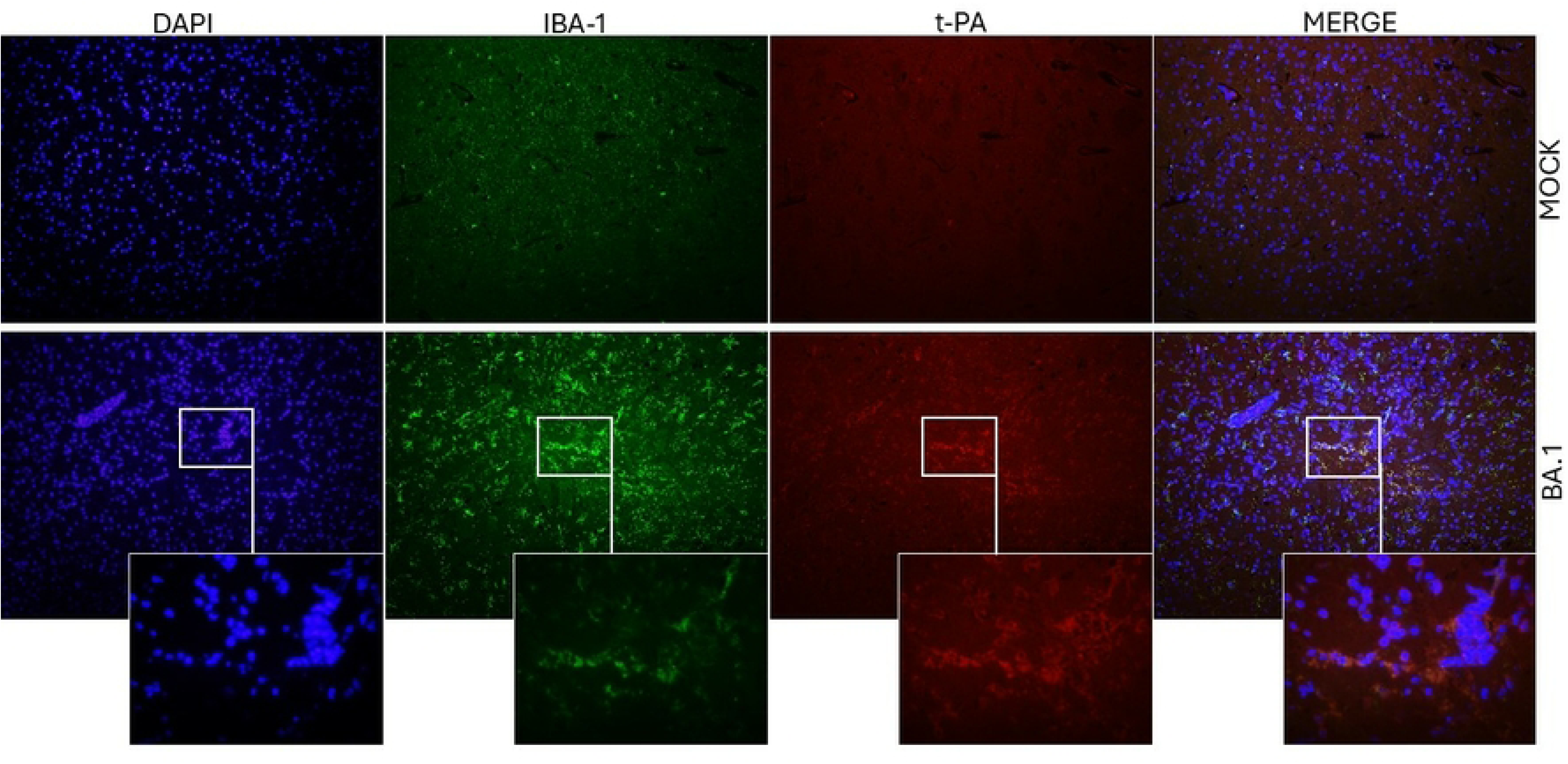
BA.1-activated microglia cells overexpress tissue-plasminogen activator (t-PA) in AC70 hACE2 Tg^+^ mice. Double-immunostaining of brain sections collected at 15 dpi. with ∼2.3×10^4^ TCID_50_ of BA.1 of SARS-CoV-2 and Mock control. t-PA (red) was co-localized with the microglia cell marker (Iba-1, green). DAPI (blue) was used to show nuclei. The bottom panels display an enlarged views of the areas marked in squares. Scale bar= 100 µm. Representative images from two independent experiments.

### BA.1-derived severe neurological symptoms were associated with disrupted blood brain barriers (BBB) integrity in AC70 hACE2-Tg mice

Among the proposed mechanisms underlying the neurological manifestations of PASC, the compromission of BBB integrity caused by SARS-CoV-2 infection is particularly attractive, as it can lead to the leakage of blood-borne components into the brain parenchyma, thereby intensifying neuroinflammation. To determine whether the disrupted BBB integrity could occur in Omicron BA.1-infected mice, we initially employed the Evans Blue dye (EBD) approach. Briefly, BA.1-infected AC70 hACE2 Tg mice were administered a 2% EBD solution via the retro-orbital (RO) route one hour before cardiac perfusion with normal saline. Mice were analyzed at the time when 1) live virus could be recovered from the lungs (∼ 7-8 dpi), 2) the onset of neurological symptoms of varying severities were identified (i.e., ∼9-10 dpi), and 3) neurological symptoms persisted in survived mice (i.e.,15 dpi). Representative photographs of the whole brains were taken, as shown in **Figure 7A-D**, to document the distinct degrees of EBD effusion. As expected, uninfected control mice did not show any tracer diffused into the brain [**Figure 7A**], confirming the intact BBB integrity. In contrast, dye leakage of varying intensities was observed in the cortex and other brain areas in 3 out of 4 mice assessed at ∼ 7-8 dpi [**Figure 7B**]. For those mice that developed the first signs of severe neurological disorders (at ∼9-10 dpi), enhanced dye extravasation could be readily observed within the cortex and cerebellum. For those mice that did not show obvious neurological abnormality at 9-10 dpi, only very weak dye leakages, if any, could be sporadically detected [**Figure 7C**]. The analysis of brains from mice with persistent neurological symptoms at 15 dpi revealed dye extravasation, particularly in the thalamus [**Figure 7D**], indicating that BBB integrity had not been restored.

**Fig 7.**
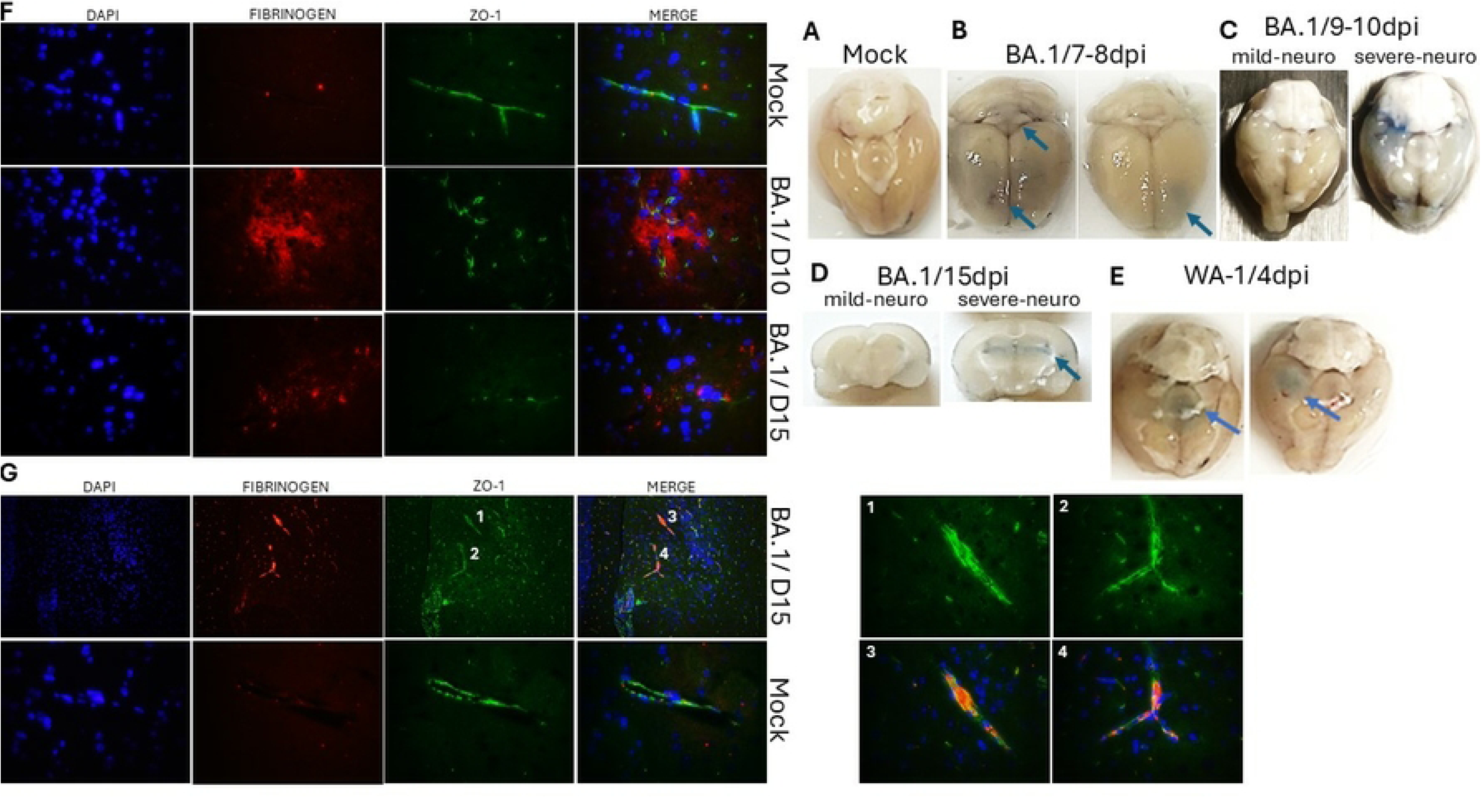
BA.1 variant of SARS-CoV-2 caused blood-brain-barrier (BBB) disruption and fibrinogen extravasation in AC70 hACE2 Tg^+^ mice. (**A-D**) AC70 hACE2 Tg mice challenged with ∼2.3×10^4^ TCID_50_ of BA.1 of SARS-CoV-2 were administered with EBD solution and cardiacally perfused with normal saline. Representative images of whole brains from (**A**) Mock, (**B**) at 7/8- (the peak of clinical disease), (**C**) at 9/10- (the onset of neurological signs), and cross-sections (**D**) at 15- dpi (the post-acute phase of infection) collected from mice with mild and severe neurological complications. (**E**) Representative images of perfused brains from mice infected with 5×10^4^ TCID_50_ of WA.1 isolate of SARS-CoV-2 at 4 dpi (the peak of clinical disease). (**F-G**) Representative double-immunostaining for fibrinogen (red) and ZO-1 (green) at the capillary level at 10 and 15 dpi. DAPI (blue) was used to label nuclei. (**F**) Infected mice exhibited extravascular fibrinogen depositions, accompanied by disrupted ZO1 expression, compared to Mock controls. **(G)** Intravascular accumulation of fibrin(ogen), accompanied by rearranged ZO-1, compared to Mock control. The right panel displays higher magnification views of infected cases.

As we were unable to recover live virus from the brains of Omicron BA.1-infected mice throughout the study, we questioned whether mice with intense viral brain infection might exhibit greater disruption of BBB integrity. Specifically, the BBB integrity in AC70 hACE2-Tg mice lethally challenged with a high-dose of WA1 isolate of SARS-CoV-2 [**26**] was examined. As shown in **Figure 7E**, the disruption of BBB at 4 dpi, when overwhelmed brain infections occurred, was unexpectedly less prominent and largely confined to the cortex, compared to those exhibited by BA.1-infected mice. Taken together, these findings imply that the extent of BBB disruption may not directly correlate with the intensity of brain infection, and other yet-to-be explored mechanisms, such as neuroinflammation, might potentially contribute to the prolonged neurological disorders.

### Neurologically affected BA.1-infected mice exhibited disrupted endothelial tight junction complexes (TJCs), fibrinogen extravasation into the brain parenchyma, and intravascular fibrin(ogen) deposits

The BBB is composed of monolayer of microvascular ECs lining the cerebral capillaries, held together with molecules tightly associated with endothelial tight junction complex (TJC). To further verify BBB disfunction, we analyzed the integrity of TJCs within the brains of mice suffered from neurological disorders. Additionally, we evaluated the content of fibrinogen within the perfused brains. Fibrinogen is not typically found in brain tissue due to the BBB restriction under physiological conditions. The perfused brain sections of Omicron BA.1-infected mice with severe neurological symptoms at ∼ 9-10 or 15 dpi were subjected to standard IF staining with specific antibodies against ZO-1, a TJC protein, and fibrinogen. As expected, neither fibrinogen deposition nor rearrangement of TJ protein ZO-1 was associated with the brain of mock-infected control. Remarkably, BA.1 infected brains showed profound extravascular fibrinogen depositions, accompanied by rearranged ZO1 expression at 9-10 dpi, with the effects persisting to some extend into the post-acute phase of infection [**Figure 7F**]. These results further imply that the leakage of plasma fibrinogen, along with other blood-borne components, into the brain of BA.1-infected mice through disrupted TJC within compromised BBB might contribute to the onset neurological disorders associated with COVID-19.

Dysregulated coagulation and fibrinolytic pathways were consistently observed within the brains of BA.1-infected AC70 hACE2 Tg mice [**Figure 5**]. Our further analysis of fibrinogen pattern revealed widespread intravascular fibrin(ogen) deposits across several brain areas, particularly in the cortex and hippocampus, where prominent vascular and microvascular blockages were observed exclusively in BA.1 infected mice [**Figure 7G**]. These findings reinforce the presence of prolonged coagulation disturbances in the CNS following BA.1 infection. Moreover, we noticed structural changes in ZO-1 within the obstructed vessels, suggesting that intravascular fibrin(ogen) buildup might be a contributing factor to TJ breakdown.

### A malfunctioning Annexin A2 (ANXA2) mechanism may contribute to the fundamental pathological aspects of PASC

Annexin A2 (ANXA2) is a key mediator of coagulation by serving as a profibrinolytic co-receptor for t-PA on endothelial cell surface, facilitating the conversion of plasminogen to plasmin. This process is fundamental for fibrinolysis and helps maintain vascular homeostasis by preventing excessive clot formation, thereby ensuring proper blood flow. Therefore, impairment of the ANXA2 mechanism may lead to a pro-coagulant state, increasing the risk of thrombosis. We have recently showed that autoantibodies targeting ANXA2 were promptly produced in mice lethally challenged with the ancestral SARS-CoV-2 WA-1 isolate, implicating their potential role in the fibrinolysis resistance phenomenon frequently reported in COVID-19 patients [**26**]. To explore whether mice infected with the less pathogenic Omicron BA.1 variant generate anti-ANXA2 autoantibodies, plasma specimens (N=5 per group) were collected at 9-10 and 21 dpi and subjected to our in-house developed ELISA analysis, as previously described [**26**]. Mock-infected plasmas were included as negative controls. As shown in **Figure 5F**, the levels of circulating anti-ANXA2 antibodies were significantly elevated at the onset of neurological manifestations (i.e. 9-10 dpi) and continued to rise by 21 dpi, with P values of 0.0021 and < 0.0001, respectively, in comparison to the uninfected control group.

ANXA2 is a multifunctional protein that facilitates the formation of actin-based adherens and tight junctions, especially through its heterotetramer (A2t) complexes, playing a pivotal role in maintaining BBB integrity. Disruption of these complexes completely aborts TJ assembly [**28**]. While we demonstrated that BA.1 infection affected TJ structures, autoantibodies targeting this protein could directly interfere with the A2t complex, leading to TJ dysfunction. To verify whether ANXA2 colocalize with TJs and the effect of BA.1 infection on ANXA2 distribution within the vessels, we performed double immunostaining of ANXA2 and ZO-1 in brain section harvested from mice infected at 9 d.p.i. and Mock controls [**Supplement Figure 2**]. We showed that in Mock mice, ANXA2 was fully colocalized with intact TJs surrounding blood vessels, whereas, in BA.1 infected mice, impaired TJ structures coexisted with diminished ANXA2 levels around the vessels. Collectively, our preliminary, albeit intriguing, results support that the ANXA2 system may play a crucial role in brain vascular homeostasis, and its dysregulation by specific autoantibodies induced by SARS-CoV-2 infection could contribute to BBB disfunction. Additional studies are currently underway to further investigate how the dysregulated ANXA2 system may compromise the integrity of tight junction complexes (TJC) and the blood-brain barrier (BBB), aiming to enhance our understanding of neuro-COVID-19 pathogenesis.

## Discussion

This study showed that the less pathogenic BA.1 Omicron variant of SARS-CoV-2 led to prolonged neurological sequelae in survived AC70 hACE2 Tg mice, proportional to the extent of neuroinflammation. Our results strongly suggest that disruption of BBB and endothelial TJC integrity may be a key contributor to CNS pathology, serving as a gateway for the extravasation of blood-borne components, i.e. fibrinogen, into the brain parenchyma. The sustained alterations in coagulation and fibrinolysis, leading to prominent intravascular fibrin(ogen) deposits, could be involved in this process and seem to display a distinct pattern within the CNS. Along with disrupted TJs, we also found that the expression of the ANXA2 protein was grossly dispersed throughout the vasculature. The presence of autoantibodies targeted this protein may be responsible for impaired function of ANXA2 system. As a pleiotropic protein essential for vascular homeostasis, including regulation of cell-surface fibrinolysis and maintenance of BBB integrity, dysregulation of ANXA2 mechanism may underlie key aspects of PASC. Collectively, our results indicate that AC70 hACE2 Tg mice infected with BA.1 variant provide a refined model with survivals for studying the pathophysiology of prolonged thrombotic and neurological sequelae associated with neuro- or long-COVID.

Unlike the WA-1 strain that triggered acute and uniformly lethal disease, BA.1-infected AC70 hACE2 Tg mice exhibited a more heterogeneous response, with severity varying from mild, severe to fatal in approximately 20% of cases. Milder outcomes upon BA.1 variant challenge have been reported in other permissive animal models, including several wild-type mice, i.e. 129, Balb/c and C57BL/6 mice, K18-hACE2 Tg mice, as well as hamsters [**38**]. However, we documented for the first time that BA.1 infected hACE2 Tg mice developed severe neurological sequelae following the acute phase of infection, including motor impairment and paralysis, that remained evident in roughly 27% survivors through 21 dpi, the endpoint of our study.

A growing body of research highlights neuroinflammation as a central aspect of Neuro-COVID. Our findings revealed that the severity of neurological complications positively correlated with the extend of neuroinflammatory changes, marked by prolonged mononuclear cell infiltration, along with astrogliosis and microglia activation. Neuroinflammation alone could contribute to glial cell overactivation, ultimately leading to neural circuit impairment and deterioration of neurological function [**39–41**]. The interplay between microglia and astrocytes could further escalate the neuroinflammation cascade [**42**]. Further investigation with the BA.1/AC70 hACE2 Tg mouse model is critical to assess the long-term effects of neuroinflammation on neurological dysfunction and to enhance our understanding of neuro- or long-COVID.

So far, the pathophysiological mechanism(s) underlying the neuroinflammation-associated neurological dysfunction triggered by SARS-CoV-2 remains unclear. While SARS-CoV-2 may facilitate several receptors expressed in the neurovascular unit, i.e. vimentin or neuropilin, it was proposed that direct virus invasion may be responsible for this phenomenon [**43, 44**]. However, it is challenging to show an active SARS-CoV-2 infection within the brains of confirmed COVID-19 patients. Although autopsy findings indicated the presence of viral RNA in multiply anatomic regions, including brain, for up to 230 days since the onset of diseases caused by the early virus strain, a direct linkage between persistent infection and the prolonged neurological sequelae has yet to be established [**45**]. While viral infection is still considered a likely trigger for neurological symptoms in COVID-19 patients, recent studies have shown that the accumulation of SARS-CoV-2 spike (S) protein in both humans and mice could induce dysregulated inflammatory responses and neuropathological changes, suggesting that persistent neurotoxic S protein at the brain borders may contribute to lasting neurological sequelae of COVID-19 [**46**]. Despite the failure to isolate infectious BA.1 viral particles from the brain tissues, we were able to detect low levels of viral RNA and S protein in selected brain regions of infected AC70 hACE2 Tg mice, suggesting either a low grade of active viral infection or accumulation of S protein or other viral components. However, given that viral spike protein was predominantly present during the peak of infection and gradually decreased afterward, it is difficult to fully claim that direct viral involvement is the primary driver of progressive inflammation and persistent neurological symptoms in infected mice.

Among other mechanisms that could contribute to or even drive the neurological sequelae after SARS-CoV-2 infection, the disruption of BBB integrity has been notably implicated. BBB is a dynamic physical and metabolic barrier primarily composed of specialized eECs, along with other resident cells, that maintains brain homeostasis by strictly regulating the influx of blood-borne molecules from the bloodstream to the CNS, thereby supporting its proper function [**47, 48**]. In various brain dysfunctions, including Traumatic Brain Injury (TBI), the compromise of the BBB is commonly associated with secondary effects such as edema, inflammation, and hyperexcitability [**49**]. In this regard, we also showed a positive correlation between the severity of neurological sequelae and the degree of BBB impairment in surviving hACE2 Tg mice infected with BA.1. We documented signs of BBB disfunction as early as 7 dpi, approximately two days prior to the onset of visible neurological symptoms, which progressively worsened and persisted through 9 to 15 dpi in infected mice with severe neurological impairments. In line with our findings, the association between neurological dysfunction and BBB breakdown has also been recently observed in patients during acute infection and in those with long-COVID brain fog symptoms [**25**].

TJs between neighboring endothelial cells form the structural basis of the BBB, acting as gatekeepers for the paracellular passage of blood-derived substances. Our data reveal that substantial TJ structural alterations coincided with fibrinogen leakage into the brain parenchyma during the active phase of infection and continued through recovery, supporting the association between BBB leakage and TJ impairment. Fibrinogen, beyond its well-established involvement in coagulation and vascular diseases, has been widely recognized for its proinflammatory potential in neurological disorders, including Alzheimer’s disease, multiple sclerosis, and traumatic brain injury (TBI) [**50–54**], stem from its ability to directly interact with neural receptors and regulatory proteins [**55**].

Insights from other animal model, studies on K18 Tg mice lethally infected with the ancestral WA-1 strain, revealed IgG leakage and TJ weakening, along with high live virus yields in the brain [**56**]. We also showed that AC70 hACE2 Tg mice lethally challenged with WA-1 exhibited a high virus level in the brain, yet the extent of BBB disruption was significantly lower than that caused by the less pathogenic BA.1 variant. Compared to BA.1 variant, the ancestral strain elicits only mild neuroinflammatory responses [**26**], likely due to an immunosuppressive effect caused by profound viral replication. Thus, the ability of infected hosts to elicit immune responses likely play key roles in shaping disease severity and its ultimate outcomes.

Our data indicates persistently elevated D-dimer levels in circulation of BA.1-infected AC70 hACE2 Tg mice, suggesting systemic prothrombotic state beyond the acute infection phase. High plasma D-dimer is a hallmark of severe COVID-19 pneumonia associated with poor outcomes. Notably, clinical studies have also reported that elevated D-dimers remain elevated during convalescence and in long-COVID cases [**57, 58**], suggesting that hypercoagulability extends beyond the acute phase and may contribute to long-term sequelae, including neurological impairments. Recent transcriptomic data from cognitively impaired patients further emphasized dysregulated coagulation pathway as a key factor [**25**]. However, the interplay between coagulation disturbances and neurological impairment in disease progression remains elusive. Our findings support the notion that coagulation disorders contribute to neuro-COVID, and further imply that thrombotic events evolve dynamically, arising locally with disease progression in an organ-specific manner rather than being merely a residual effect of acute infection. We noted that the levels of tissue-associated D-dimer expression fluctuated in a tissue/organ-specific manner in BA.1-infected AC70 hACE2 Tg mice, corresponding with disease progression. Specifically, we showed that lungs were the primary early target of SARS-CoV-2 infection, with host immune responses typically peaking at 7 dpi, closely aligning with increased D-dimer levels and intravascular clot formation. D-dimers are the major fibrinolysis-specific degradation products, the main component of insoluble clot, released from crosslinked fibrin within injured tissue by the action of plasmin, the active form of plasminogen, in the presence of t-PA. Detection of d-dimer antigen indicates enhanced fibrinolysis. In the lungs of infected mice, this process appears to be transient, as histopathological examination at later stages showed no signs of infiltrating mononuclear cells or clot formation. In contrast, brain pathology typically emerged at 7 d.p.i., with the neuroinflammatory response intensifying by 21 d.p.i., corresponding with rising D-dimer levels and suggesting ongoing or progressive coagulation disturbances. Moreover, analysis of t-PA expression, the key enzyme in fibrinolysis, in both organs revealed that the fibrinolytic pathway was predominantly overactivated in the brain, suggesting a distinct pattern of fibrinolysis in the CNS. Endothelial cells within the microvasculature have been recognized as the main source of t-PA, and its function has been conventionally assigned to fibrinolysis [**34, 35**]. However, existing research, particularly in mice lacking specific fibrinolytic system components, has detected t-PA in brain cells, including glial cells and neurons [**36, 59, 60**], extending its function from thrombolysis to involvement in CNS physiology and pathology, including learning and memory, anxiety, epilepsy, stroke, Alzheimer’s disease, or spinal cord injury [**61–66**]. Our findings show a positive correlation between t-PA expression and neuroinflammation progression. Additionally, its close association with Iba-1-positive cells suggests that activated microglia cells could contribute to t-PA secretion in response to BA.1 infection. Given the intricate role of the t-PA/plasminogen system in the CNS, further clarification of its contributions could lead to new insights into the mechanism of neuro- or long-COVID.

Interestingly, despite t-PA overexpression, the prolonged presence of intravascular fibrin(ogen) deposits in the infected brain of AC70 hACE2 Tg mice indicates incomplete fibrinolysis. Many studies have documented thrombosis in COVID-19 patients, even with anticoagulant treatment [**67, 68**], or increased fibrinolysis markers, strongly suggesting fibrinolysis resistance. We have recently reported that WA-1 infection led to the rapid production of autoantibodies targeting Annexin A2 (ANXA2), which could interfere with the ANXA2-S100A10 association in AIIt complexes on the cell surface, ultimately impairing t-PA-dependent plasmin formation [**26**]. While we demonstrated that infection with BA.1 variant also triggered anti- ANXA2 autoantibody production that enhanced over the course of the infection, the dysregulation of the ANXA2 system may have been a contributing factor to fibrinolysis failure in the post-acute phase of the disease.

Beyond the role played in fibrinolysis process, studies using ANXA2 knockout (A2KO) mouse model demonstrated that ANXA2 contributes to regulation of the cerebral endothelial integrity under physiological and pathological conditions, suggesting that ANXA2 system may play a pivotal role in BBB development, and in protection of its functional and structural integrity after brain injury [**69, 70**]. Moreover, exogenous administration of recombinant ANXA2 noticeably improved BBB integrity and upregulated BBB junction expression in TBI mouse model [**71**]. In line with these findings, we showed that ANXA2 was markedly disorganized in the brains of BA.1-infected mice with compromised TJs, whereas it remained intact in uninfected mice. AIIt complexes exhibit greater bundling activity compared to ANXA2 monomers [**72**], and the addition of a synthetic peptide that interferes with ANXA2-S100A10 binding completely abolishes TJ assembly [**28**]. Thus, ANXA2, particularly in its AIIt complex form, appears to be a crucial membrane protein that ensure CNS hemostasis by regulating TJ integrity, and the presence of anti-ANXA2 autoantibodies may interfere with this process. Studies are ongoing to further elucidate the contribution of the ANXA2 system to neuro-COVID.

In summary, we have established that hACE2 Tg mice infected with the BA.1 variant of SARS-CoV-2 provide a distinctive platform for studying neuro-COVID and PASC. Lasting BA.1-induced neurological effects were associated with the degree of BBB disruption, neuroinflammatory responses and coagulation disorders. Our findings indicate that coagulation and fibrinolysis pathways exhibit a distinct pattern in the brain, with their dysregulation potentially driven, at least in part, by glial cell activation, suggesting that local coagulopathic changes may stem from ongoing neuroinflammation. Conversely, BBB dysfunction may facilitate the entry of plasma components like fibrinogen, into the brain parenchyma, potentially initiating coagulopathic processes. Furthermore, a systemic prothrombotic state may promote fibrin(ogen) accumulation at vascular walls, leading to TJ disruption and BBB impairment. These insights reveal a dynamic relationship between the coagulation/fibrinolysis system, neurovascular integrity, and neuroinflammatory mechanisms. Lastly, we extended our exploration of the potential role of ANXA2 system in these processes. Further research into ANXA2 involvement in cerebrovascular homeostasis could lead to innovative strategies for addressing neuro-COVID.

## Materials and Methods

### Ethics statement

All experiments involving animals and infectious viruses were conducted at the Galveston National Laboratory (GNL) at The University of Texas Medical Branch at Galveston (UTMB), Texas, an AAALAC accredited (November 24, 2020) and PHS OLAW approved (February 26, 2021) high-containment National Laboratory. All procedures followed protocols approved by the Institutional Animal Care and Use Committee (IACUC) at UTMB Galveston.

### Viral stock and cell culture

SARS-CoV-2 (BA.1 Omicron variant, strain: EHC_C19_2811c), obtained from the World Reference Center for Emerging Viruses and Arboviruses (WRCEVA), was used throughout the studies. Viral stock was propagated in Vero E6 cells (American Type Culture Collection) and the resulting cell-free supernatants were subsequently sequenced, titrated, and stored at – 80°C as working stocks. For all studies, the maximum achieved viral dose, roughly 2.3×10⁴ TCID₅₀, was consistently used.

### Mice challenge and health monitoring

Human ACE2 transgenic mice, line AC70 (AC70 hACE2 Tg), were obtained from Taconic Biosciences (NY, Cat # 18222). Six-to-eight-week-old mice were anaesthetized with isoflurane in oxygen and inoculated intranasally (IN) with BA.1 variant of SARS-CoV-2 in 60 μl of Eagle’s minimum essential medium (EMEM) supplemented with 2% heat-inactivated Fetal Bovine Serum (FBS) (M-2). Infected animals were monitored at least once daily for weight changes, other signs of illness, and mortality.

### Tissue and blood collection

Blood samples were collected via retroorbital (RO) route under anesthesia. At the designed time point post-infection (p.i.) mice were euthanized via CO2 inhalation, and lung and brain tissue specimens were harvested. Briefly, the right lung was flushed with Phosphate Buffered Saline (PBS) and subsequently inflated with 10% neutral buffered formalin for histopathology and immunohistochemistry. The left lung and brain tissues were divided for downstream experiments, including viral titration, RNA and protein extraction. For certain experiments, mice were subjected to cardiac perfusion before proceeding with the next steps. Briefly, anesthetized mice were subjected to a slow infusion of 30 ml of normal saline via the left ventricle using a 26 G needle, until the fluid draining from the right atrium became clear.

### Blood-Brain-Barrier (BBB) assessment using Evans Blue (EB) assay

A 2% solution of EB dye (Sigma) in 0.9% normal saline (4 mL/kg of body weight) was injected via RO route at designed points p.i. The dye was allowed to circulate for an hour, and then the mice were transcardially perfused according to the aforementioned method. The brains were harvested, and representative pictures were taken.

### Virus titration

To quantify infectious virus, the collected tissues were weighed and homogenized in PBS containing 10% FBS using the TissueLyser (Qiagen). The supernatants were clarified by centrifugation and then titrated in Vero E6 cell cultures in 96-well microtiter plates using a standard infectivity assay [**Tseng 2007 32**]. Viral titers were expressed as TCID_50_/g of tissue (log_10_).

### Histopathology (H&E), immunohistochemistry (IHC) and immunofluorescence (IF)

For H&E staining, tissue samples were fixed in 10% neutral buffered formalin (Sigma) for 72 hours, followed by transfer to 70% ethanol and paraffin embedding. Staining on deparaffinized sections was performed by using routine hematoxylin-and-eosin method. IHC for evaluating microglia activation was performed according to the previously established protocol [**32**] using anti-IBA1 rabbit monoclonal antibodies (<u>GeneTex,</u> Cat# GTX637629) at 1:500 dilution. For IF staining, the slides were incubated with primary antibodies: anti-SARS-CoV-2 S glycoprotein rabbit polyclonal antibody (Abcam, Cat# ab272504, 1:1000), anti-GFAP chicken polyclonal antibody (GeneTex, Cat# GTX85454, 1:500), anti-IBA1 mouse monoclonal antibody (Abcam, Cat# ab283319, 1:100), anti-TPA rabbit monoclonal antibody (Abcam, Cat# ab157469, 1:100), anti-ZO1 mouse monoclonal antibody (Invitrogen, Cat# 33-9100, 1:100) or anti-ANXA2 rabbit polyclonal antibody (Abcam, Cat# ab41802, 1:100) diluted in background-reducing antibody diluent (Dako, Cat# S302283-2). Next, the slides were incubated with secondary antibodies: goat anti-rabbit IgG Alexa Fluor 568 (Invitrogen, Cat# A-11011), goat anti-chicken IgY Alexa Fluor 488 (Invitrogen, Cat# A-11039), or goat anti-mouse IgG Alexa Fluor 488 (Invitrogen, Cat# A-1101), all diluted at 1:1000. Coverslips were then mounted using ProLong Gold antifade reagent with DAPI (Invitrogen, Cat# P36931). For fibrin/ fibrinogen staining, slides were blocked with 5% BSA/ 0.3% Triton X-100 in PBS, followed by incubation with polyclonal rabbit anti-human fibrinogen (DAKO, Cat# A0080) at a 1:1000 dilution in 1% BSA/0.1% Triton X-100 in PBS. Immunofluorescence microscopy (Olympus DP71) was used for visualization.

### ELISA assay

Gamma-irradiated EDTA-plasma and tissue specimens collected from infected and uninfected mice were used for ELISA assay. D-Dimer (D2D) levels were measured using a commercially available kit (ABclonal, Cat# RK02735), and absorbance was recorded with an ELISA plate reader (Molecular Devices). The levels of anti-ANXA2 antibodies were assessed following a previously published in-house protocol [**26**].

### Western Blot (WB)

Gamma-irradiated tissue specimens collected from infected and uninfected mice were homogenized on ice in RIPA buffer (Abcam, Cat# ab65400) using Dounce Glass Manual Tissue Grinder (Etcon). The total protein concentration was quantified using Pierce BCA Protein Assay Kit (Thermo Fisher). Samples were then separated by gel electrophoresis followed by immunoblotting. Equal amounts of proteins were subjected to 4–20% SDS-polyacrylamide gel (Bio-Rad) electrophoresis (SDS-PAGE). Proteins were transferred onto a polyvinylidene difluoride membrane and then incubated with anti-TPA rabbit monoclonal antibodies (Abcam, Cat# ab157469, 1:1000), followed by incubation with Eu-Labeled Goat Anti-Rabbit ScanLater (Molecular Devices) for an hour. Blots were visualized using ScanLater Western Blot Detection System (Molecular Devices).

### RNA extraction and real-time quantitative reverse qRT-PCR

Pieces of tissues were stored in RNAlater solution (Qiagen) and then processed for homogenization as outlined above. Total RNA was extracted using TRIzol (Life Technologies), treated with Turbo DNAse I (Invitrogen), and quantitated using SpectraMax Paradigm Microplate Reader (Molecular Device). RT-qPCR was performed using the SuperSAcript III One-Step RT-PCR System with Platinum Taq DNA Polymerase (Invitrogen). The sequences for the primers and probe used to amplify the SARS-CoV-2 envelope (E) gene were as follows: forward primer: 5′ACAGGTACGTTAATAGTTAATAGCGT-3′; reverse primer: 5′ATATTGCAGCAGTACGCACACA-3′; probe: FAM-ACACTAGCCATCCTTACTGCGCTTCG-BBQ1. For quantitation of viral RNA, a standard curve was generated using the 2019-nCoV_N Positive Control plasmid (Integrated DNA Technologies, Cat# 10006621).

### Statistical Analysis

Statistical analyses were conducted using GraphPad Prism software. Data from multiple experiments were presented as mean ± standard error of the mean (SEM). For multiple group comparisons, one-way analysis of variance (ANOVA) was applied, while Student’s t-test was used for pairwise comparisons. A two-tailed P value of < 0.05 was considered statistically significant.

## Acknowledgments

We appreciate the support of the Animal Resources Center for their logistical assistance and dedicated animal care, and we thank the Research Histology Core for their tissue processing expertise. We thank Dr. Kenneth S. Plante for providing the virus strain, and the World Reference Center for Emerging Viruses and Arboviruses (WRCEVA), Dr. Natalie Thornburg and the Centers for Disease Control and Prevention (CDC) for graciously donating the virus to the collection.

## Supporting information

**S1 Fig. The comparison of neuropathology in AC70 hACE2 Tg^+^ mice that developed mild versus severe neurological manifestations at post-acute phase of disease.** Brain tissue specimens harvested at 14- and 21-d.p.i. from AC70 hACE2 Tg^+^ mice challenged with ∼2.3×10^4^ TCID_50_ of BA.1 of SARS-CoV-2 were subjected to the histopathology study. Briefly, paraffin-embedded, and formalin-fixed tissue sections were stained with hematoxylin and eosin (H&E), as described in Materials and Methods. The comparative examination between severe (**B&D**) and mild (**A&C**) neuro-cases. At 14 d.p.i., more extensive inflammation and perivascular cuffing with microglial activation was observed in mice with severe (**B**) compared to mild (**A**) neuro-cases. At 21 d.p.i, persistent neuroinflammation observed in severe cases (**D**). No detectable pathology was seen in the brain (**C**) of mouse without severe neurologic symptom. Scale bar = 80µm. Representative images from two independent experiments.

**S2 Fig. BA.1 variant of SARS-CoV-2 affected Annexin A2 (ANXA) distribution within blood vessels in AC70 hACE2 Tg^+^ mice.** Double-immunostaining of brain sections collected at 10 d.p.i. with ∼2.3×10^4^ TCID_50_ of BA.1 of SARS-CoV-2, and Mock control. ANXA2 (red) was co-localized with the TJ marker (ZO-1, green). DAPI (blue) was used to show nuclei.

